# Identification and mechanistic analysis of an inhibitor of the CorC Mg^2+^ transporter

**DOI:** 10.1101/2021.02.10.430528

**Authors:** Yichen Huang, Kaijie Mu, Xinyu Teng, Yimeng Zhao, Yosuke Funato, Hiroaki Miki, Weiliang Zhu, Zhijian Xu, Motoyuki Hattori

## Abstract

The CorC/CNNM family of Na^+^-dependent Mg^2+^ transporters is ubiquitously conserved from bacteria to humans. CorC, the bacterial member of the CorC/CNNM family of proteins, is involved in resistance to antibiotic exposure and in the survival of pathogenic microorganisms in their host environment. The CorC/CNNM family proteins possess a cytoplasmic region containing the regulatory ATP-binding site. While CorC and CNNM have attracted interest as therapeutic targets, inhibitors targeting the ir regulatory ATP-binding site have not yet been identified.

Here, we performed a virtual screening of CorC by targeting its regulatory ATP-binding site, identified a chemical compound named IGN95a with inhibitory effects on both ATP binding and Mg^2+^ export, and determined the cytoplasmic domain structure in complex with IGN95a. Furthermore, a chemical cross-linking experiment indicated that with ATP bound to the cytoplasmic domain, the conformational equilibrium of CorC was shifted more towards the inward-facing state of the transmembrane domain. In contrast, IGN95a did not induce such a shift. Our results provide a structural basis for the further design and optimization of chemical compounds targeting the regulatory ATP-binding site of CorC as well as mechanistic insights into how ATP and chemical compounds modulate the transport activity of CorC.

## Introduction

CorC, a prokaryotic member of the CorC/CNNM family of proteins, is involved in Mg^2+^ transport^1–5^. In the pathogenic bacterium *Staphylococcus aureus*, CorC confers resistance to the high concentrations of Mg^2+^ ions in the infected host, increasing the pathogenicity of the bacterium^3,5^. Upon exposure to ribosome-targeting antibiotics, the expression of CorC is upregulated in the L22* strain of *Bacillus subtilis* to enhance Mg^2+^ flux for resistance to antibiotics^4^. Furthermore, in humans, CNNM proteins, eukaryotic members of the CorC/CNNM family of proteins, are involved in a number of biological events, such as body absorption/reabsorption of Mg^2+^, hypertension, genetic disorders and tumour progression^6–13^. Therefore, CorC and CNNM are possible targets for novel antibiotics and drugs for treating various diseases, such as cancer.

CorC/CNNM family proteins share a conserved transmembrane (TM) DUF21 domain and a cytoplasmic cystathionine-beta-synthase (CBS) domain with the regulatory ATP-binding motif^1,3,14,15^. The recently determined structures of the TM and CBS domains of CorC from *Thermus parvatiensis* (TpCorC) revealed the mechanisms of Mg^2+^ and ATP binding, respectively^16^. Furthermore, multiple structures containing the CBS domain of CNNM proteins have also been reported thus far^17–24^.

While the Na^+^ gradient is implicated as a driving force for Mg^2+^ export from CorC and CNNM^6,16^, ATP binding to the CBS domain of CorC and CNNM proteins is essential for the regulation of Mg^2+^ efflux activities^16,24,25^. Therefore, chemical compounds targeting the CBS domain of the CorC/CNNM family proteins, especially their ATP-binding site, could be exploited for therapeutic interventions against various diseases, but such compounds have not yet been identified, hindering the development and optimization of chemical compounds targeting the CorC and CNNM proteins. Furthermore, how ATP binding to CorC and CNNM proteins modulates their transport activity also remains unclear.

In this work, based on the ATP-bound structure of the CorC CBS domain, we performed virtual screening and functional assays to identify chemical compounds targeting the ATP-binding site of the CorC CBS domain and identified the chemical compound with inhibitory effects on ATP binding and Mg^2+^ export. Structural and biochemical analyses provided mechanistic insights for further design and optimization of chemical compounds targeting the ATP-binding site of CorC as well as mechanistic insights into how ATP and chemical compounds modulate the transport activity of CorC.

## Results

### Virtual screening

A computational strategy was applied to find potential active compounds against CorC (**Fig. 1**). First, 6412 compounds designed in-house were docked to the ATP-binding site in the CorC CBS domain structure (PDB ID: 7CFI). Then, 12 docked compounds among the compounds with the top 30 2D similarities with ligand efficiencies better than –0.40 kcal/mol, which interacted with at least three residues in the pocket via hydrogen bonds and π-π interactions, were chosen for further MM/GBSA calculations (**Fig. 2** and **Table 1**).

**Table. 1.**
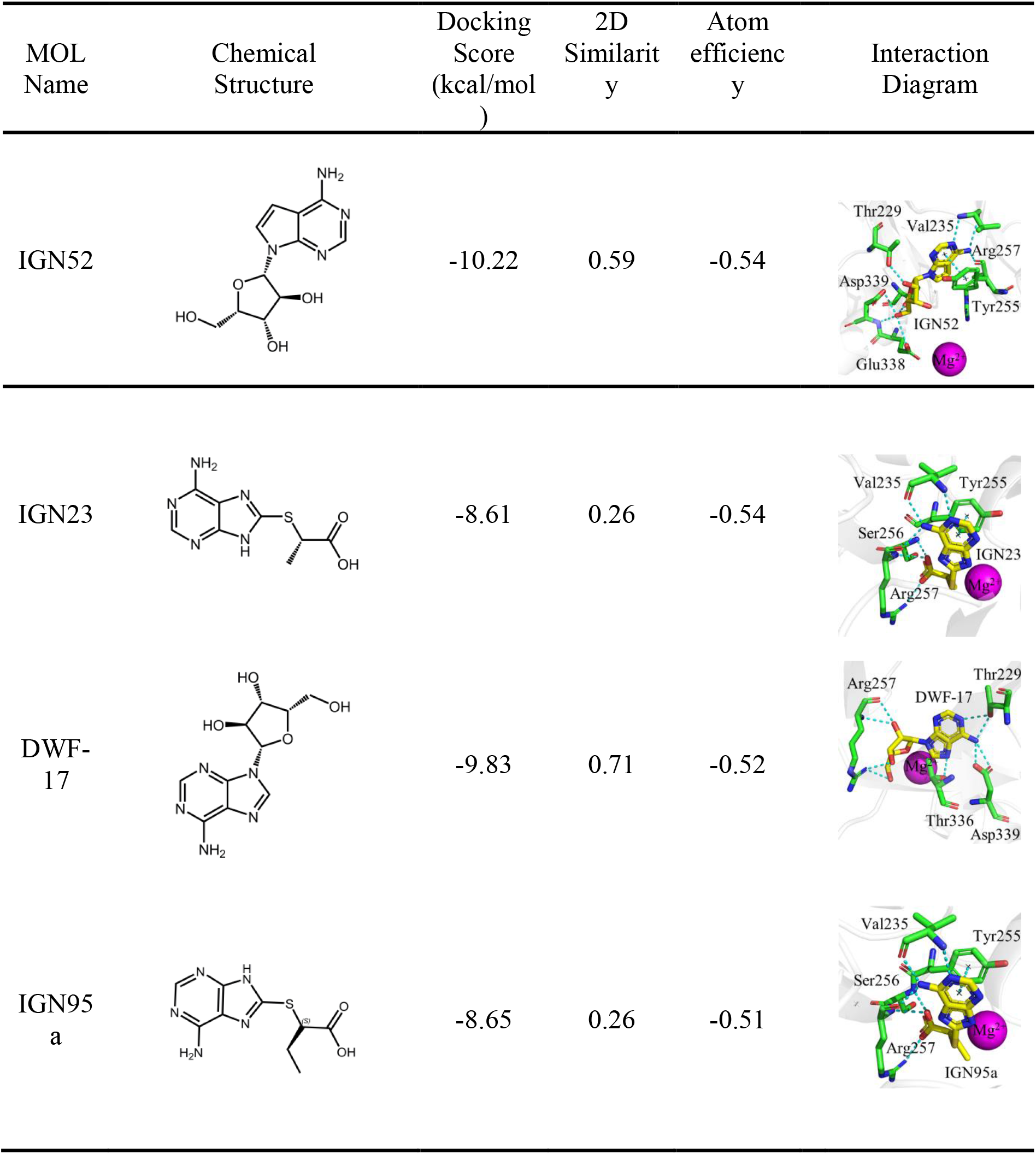
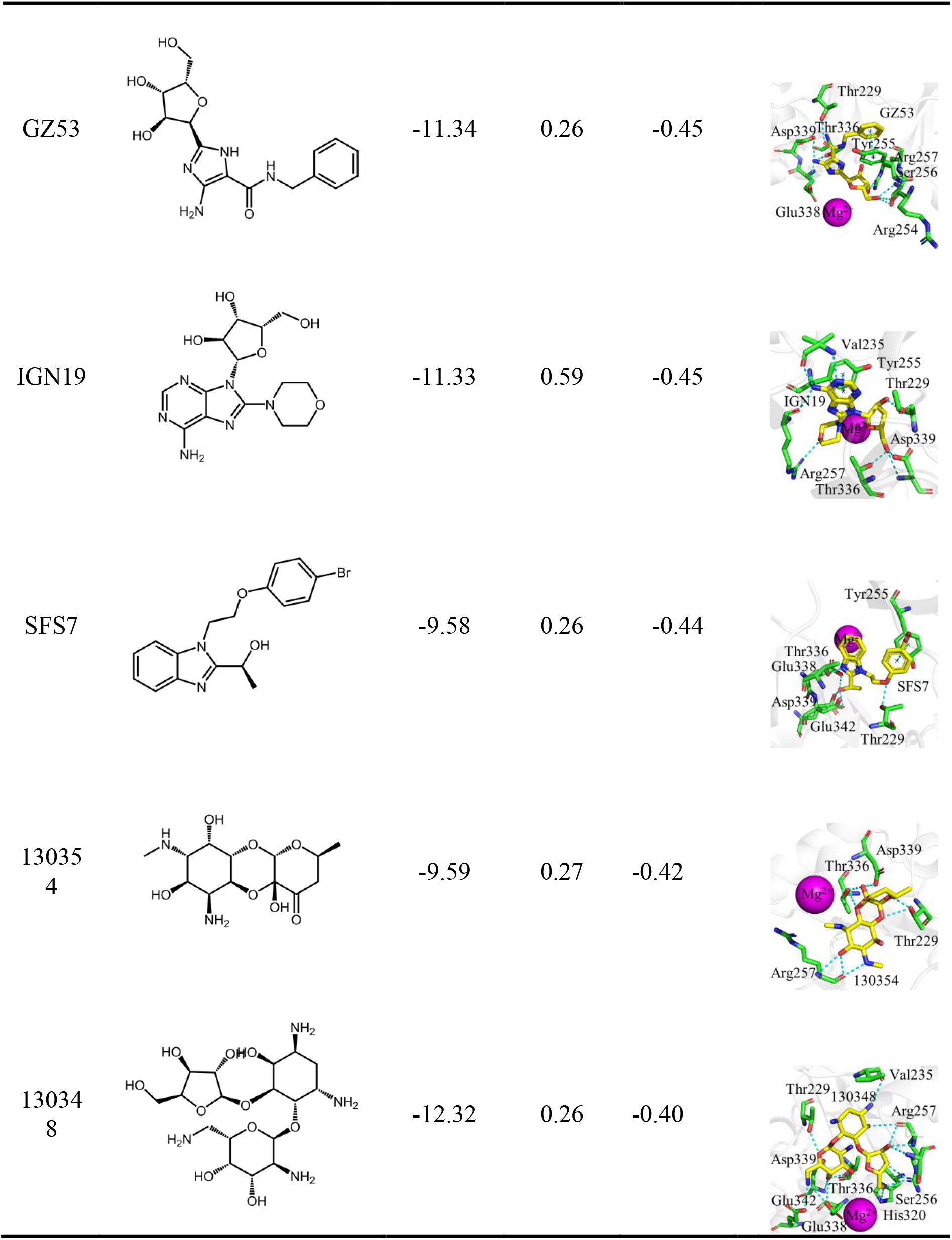
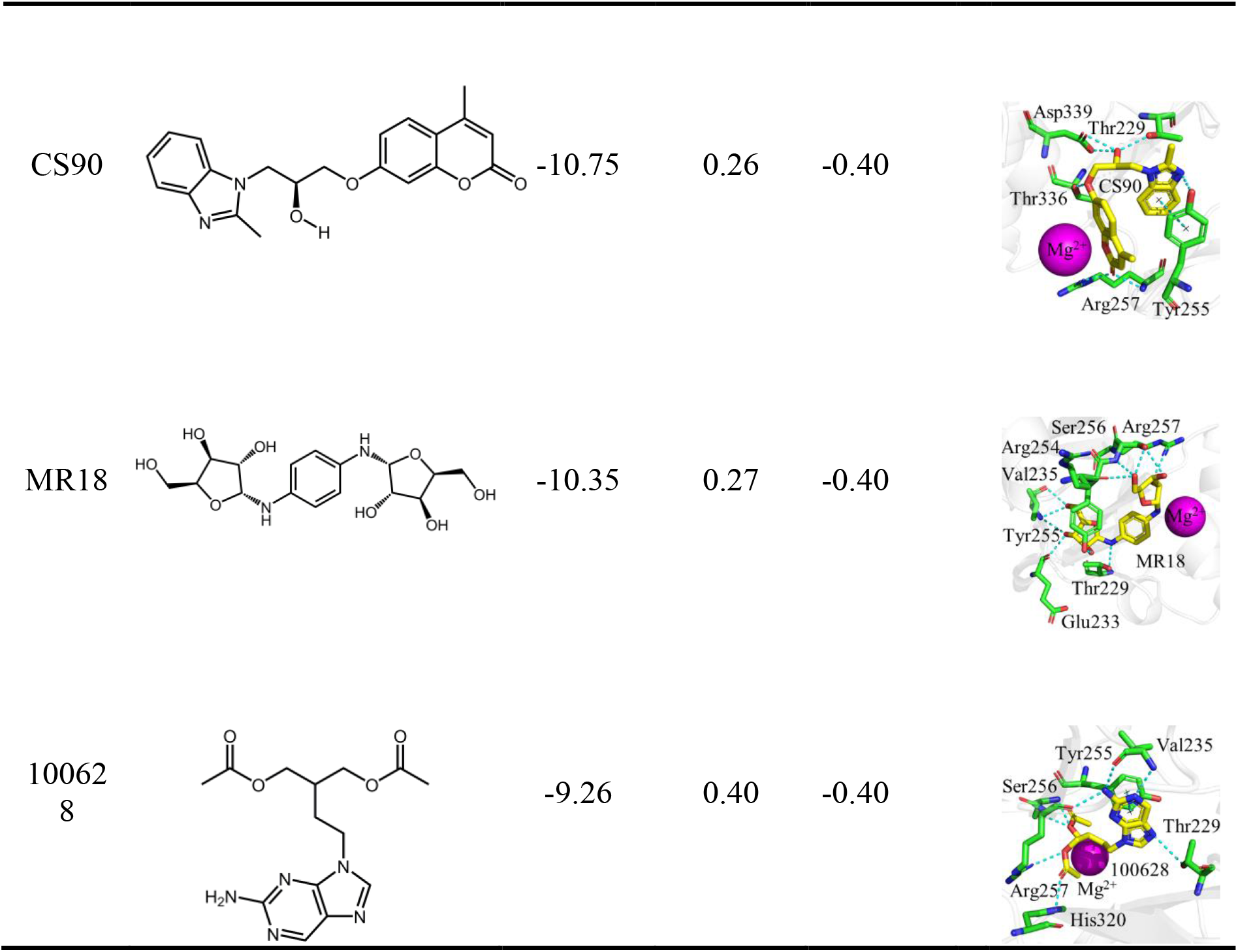
12 compounds selected are ranked by atom efficiency.

| MOL Name | Chemical Structure | Docking Score (kcal/mol) | 2D Similarity | Atom efficiency | Interaction Diagram |
| --- | --- | --- | --- | --- | --- |
| IGN52 |  | -10.22 | 0.59 | -0.54 |  |
| IGN23 |  | -8.61 | 0.26 | -0.54 |  |
| DWF-17 |  | -9.83 | 0.71 | -0.52 |  |
| IGN95a |  | -8.65 | 0.26 | -0.51 |  |
| GZ53 |  | -11.34 | 0.26 | -0.45 |  |
| IGN19 |  | -11.33 | 0.59 | -0.45 |  |
| SFS7 |  | -9.58 | 0.26 | -0.44 |  |
| 13035<br>4 |  | -9.59 | 0.27 | -0.42 |  |
| 13034<br>8 |  | -12.32 | 0.26 | -0.40 |  |
| CS90 |  | -10.75 | 0.26 | -0.40 |  |
| MR18 |  | -10.35 | 0.27 | -0.40 |  |
| 10062<br>8 |  | -9.26 | 0.40 | -0.40 |  |

**Fig. 1.**
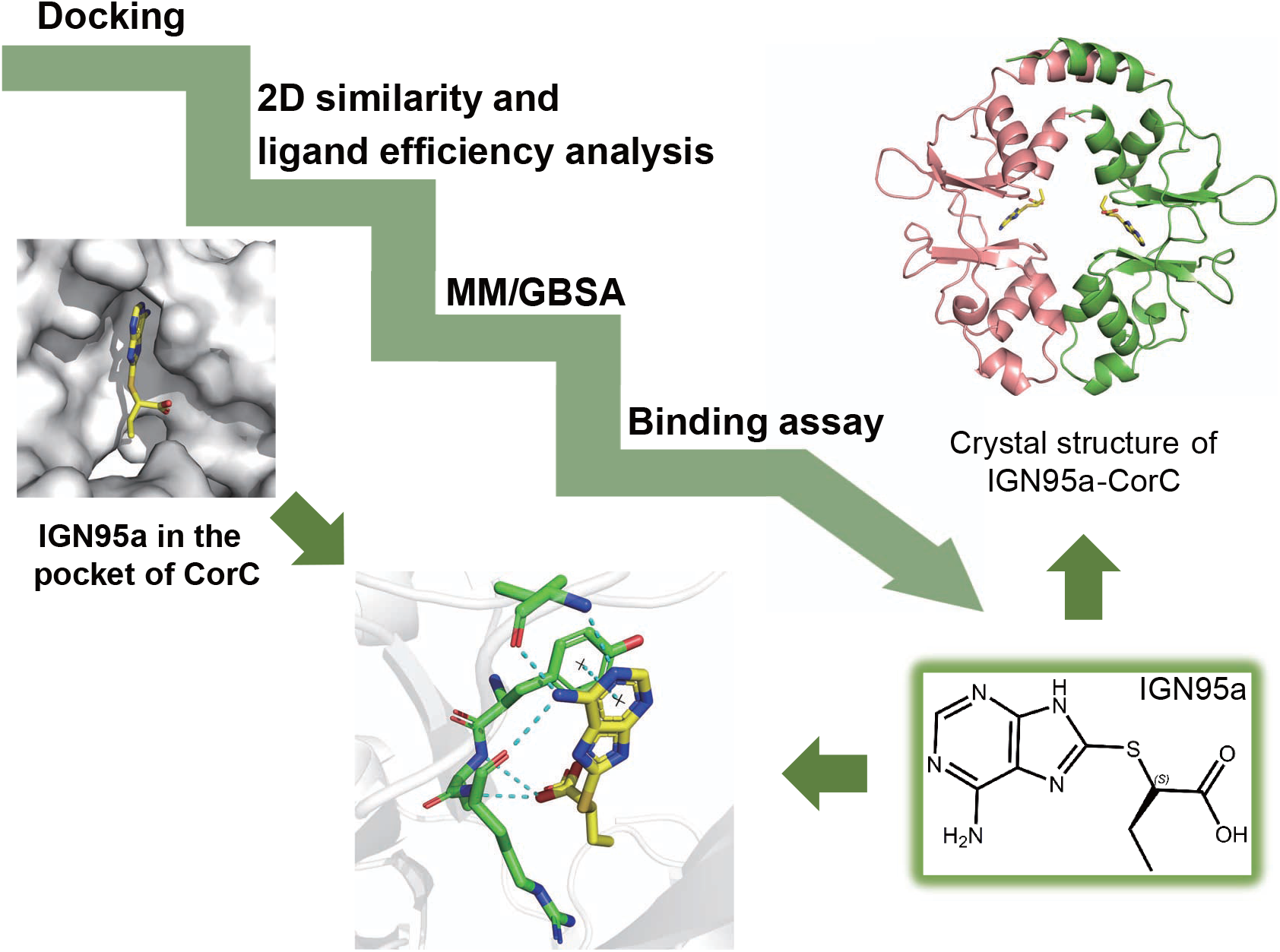
Computational strategy to identify active compounds. Molecular docking, 2D similarity analysis, atom efficiency analysis, and MM/GBSA calculations were combined to identify active compounds against CorC.

**Fig. 2.**
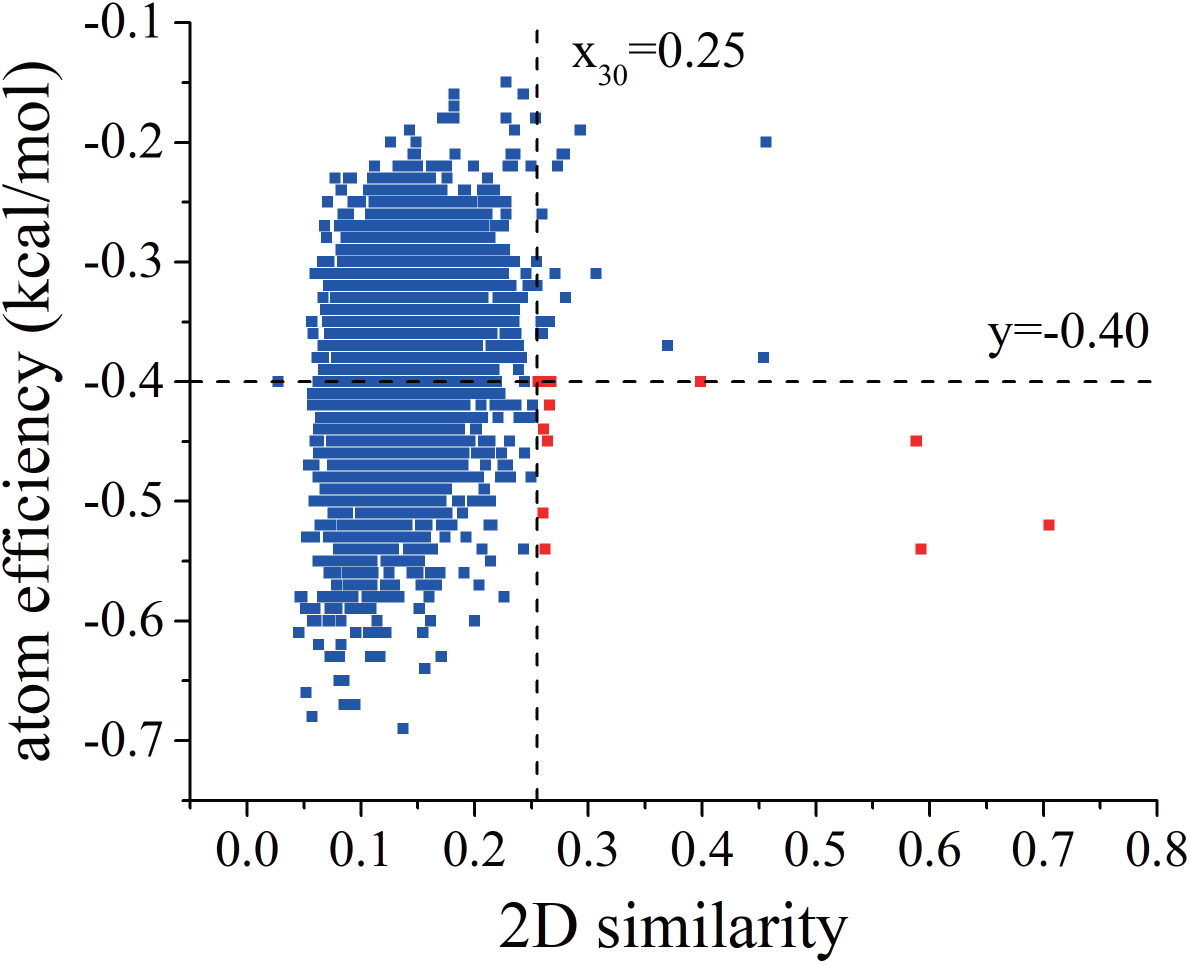
Compound selection from 2D similarity and atom efficiency analysis. The top 30 compounds with the highest 2D similarity to ATP were selected first. Among them, 12 compounds with atom efficiencies better than −0.40 (coloured red) were chosen for further MM/GBSA calculations.

To identify the best CorC-binding compound with comprehensive considerations, the free energies of the binding of 12 compounds to CorC were calculated by the MM/GBSA method. As shown in **Table 2**, 3 available compounds (IGN95a, IGN23 and IGN19) with binding free energies greater than –10 kcal/mol were selected for the bioassay.

**Table 2.**
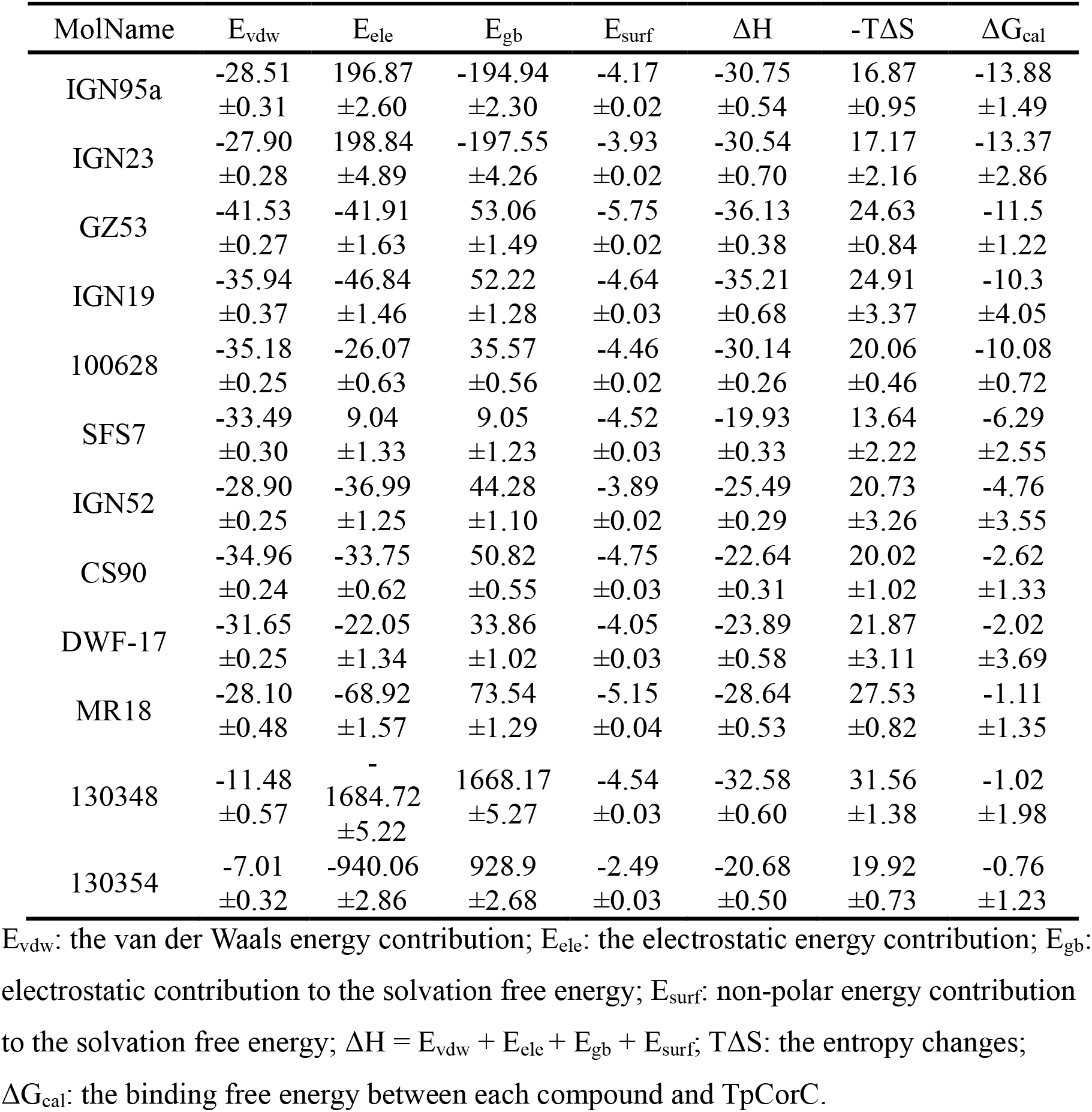
The binding free energies of 12 compounds to TpCorC calculated by MM/GBSA.

### *In vitro* screening

We then performed fluorescent ATP-based binding assays with the chemical compound candidates from the virtual screening. We employed 2′(3′)-O-(N-methylanthraniloyl)adenosine 5′-triphosphate (mant-ATP) for the binding assay and measured fluorescence resonance energy transfer (FRET) from endogenous Trp residues to the bound mant-fluorophore^26,27^.

The FRET data showed that mant-ATP was bound to the purified CorC CBS domain with a *K*_d_ of 1.36 ± 0.10 μM (**Fig. 3a**). For validation of this method, we performed a competitive binding assay with ATP (**Fig. 3b**). The addition of ATP at the respective concentrations yielded an IC_50_ value of 0.57 ± 0.08 μM, which is comparable to the previously reported *K*_d_ value of the CorC CBS domain for ATP obtained by isothermal titration calorimetry (ITC)^16^.

**Fig. 3.**
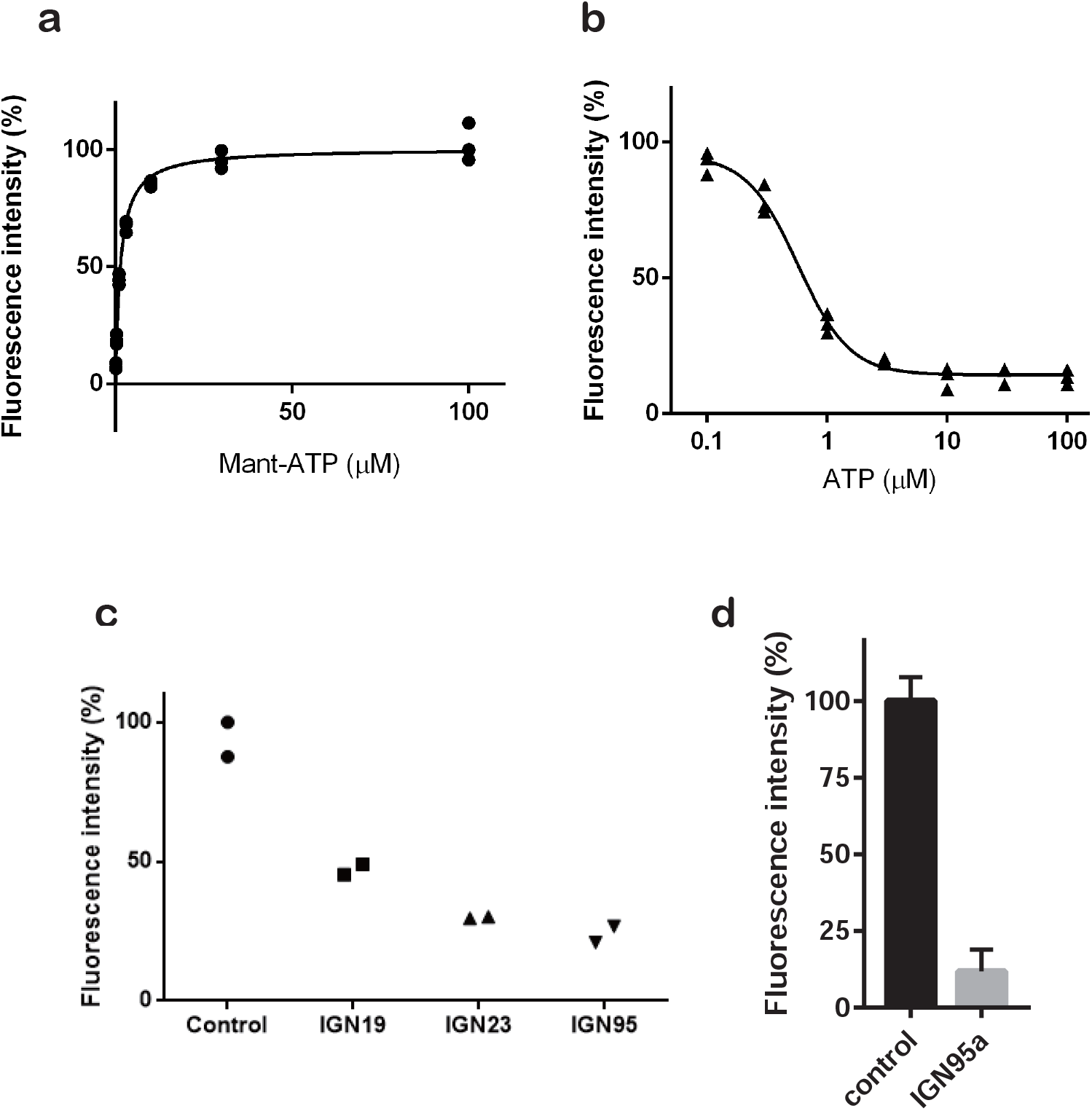
Mant-ATP binding assay. (**a**) Saturation of mant-ATP binding to the CorC CBS domain (1 μM) detected by FRET. R^2^ = 0.9989, n = 3. (**b**) ATP-based inhibition test of the binding of the CorC CBS domain (1 μM) and mant-ATP (1 μM) using FRET. R^2^ = 0.9894, n = 3. (c, **d**) Mant-ATP binding inhibition by chemical compounds. (**c**) Data are expressed as dots. n = 2. Each chemical compound was added at 0.25 mM. (**d**) Data are expressed as the mean ± SE. n = 6. IGN95a was added at 2 mM.

Finally, from the mant-ATP-based screening of chemical compound candidates from the virtual screening, we identified IGN95a, an adenine analogue, as the compound that most potently inhibited mant-ATP binding (**Fig. 3c, d**).

### Characterization of IGN95a

We further characterized IGN95a (**Fig. 4**). First, the ITC experiment confirmed IGN95a binding to the CorC CBS domain with a *K*_d_ value of 47.0 μM (**Fig. 4a and Supplementary Table 1**).

**Fig. 4.**
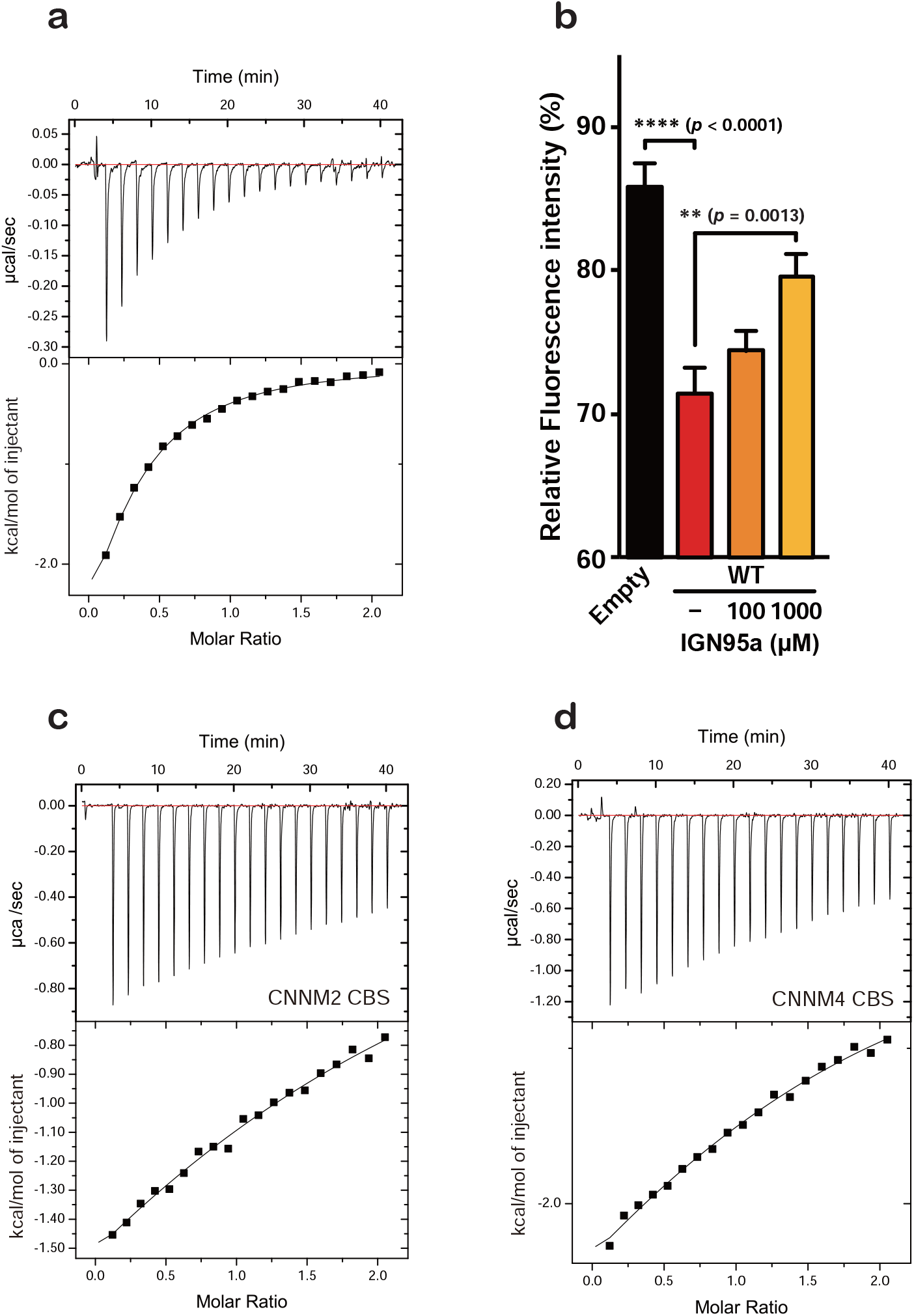
Functional characterization of IGN95a. (**a**) ITC data of the TpCorC CBS domain with IGN95a. The raw ITC data and profiles are shown. Measurements were repeated twice, and similar results were obtained. (**b**) Mg^2+^ export assay of CorC-expressing cells treated with IGN95a. Bar graph: relative fluorescence intensities after Mg^2+^ depletion (at 5 minutes, mean ± SEM, Empty: n = 10, WT without IGN95a: n = 18, WT with 100 μM IGN95a: n = 19, and WT with 1000 μM IGN95a: n = 30). (**c, d**) ITC data of the CBS domain of CNNM2 (**c**) and CNNM4 (**d**) with IGN95a.

We then tested the effects of IGN95a on the Mg^2+^ export activity of CorC (**Fig. 4b**). For the Mg^2+^ export activity assay with Magnesium Green, a fluorescent indicator dye for Mg^2+^, we employed the human embryonic kidney 293 (HEK293) cell line stably expressing CorC at the cell surface, as it was employed for the previous structure-based mutational analysis of CorC^16^. While the intensity of the fluorescent signal in the control cells expressing CorC soaked with a buffer containing only 1% DMSO decreased after the removal of Mg^2+^ ions from the bath solution, we observed inhibitory effects of IGN95a on the Mg^2+^ export activity of CorC after soaking with IGN95a in 1% DMSO buffer with HEK293 cells expressing CorC (**Fig. 4b**). These results show that IGN95a acts as an inhibitor of both ATP binding and Mg^2+^ export.

Furthermore, we tested IGN95a binding to the CBS domain of human CNNM2 and CNNM4 (**Fig. 4c, d)**. In the ITC experiments, IGN95a showed weak binding to CNNM2 and CNNM4. The exact *K*_d_ values could not be estimated because the titrations were not completed (>1000 μM for the CBS domain of CNNM2, > 500 μM for the CBS domain of CNNM4). These results suggest the specificity of IGN95a against CorC compared to CNNM family proteins, which may be beneficial for future optimization.

### Inhibitor-bound structure

To understand the molecular interactions between the CorC CBS domain and IGN95a, we performed co-crystallization of the TpCorC CBS domain with IGN95a. Since the wild-type CBS domain protein has a very high affinity for ATP, it was difficult to completely remove endogenous ATP during purification, which is not ideal for co-crystallization with IGN95a. Therefore, we employed the T336I mutant with weaker affinity for ATP^16^ because mutation at Thr336 was relatively unlikely to affect IGN95a binding based on the initial docking model. Indeed, we successfully determined the crystal structure of the TpCorC CBS domain in complex with IGN95a (**Fig. 5a and Supplementary Fig. 1**).

**Fig. 5.**
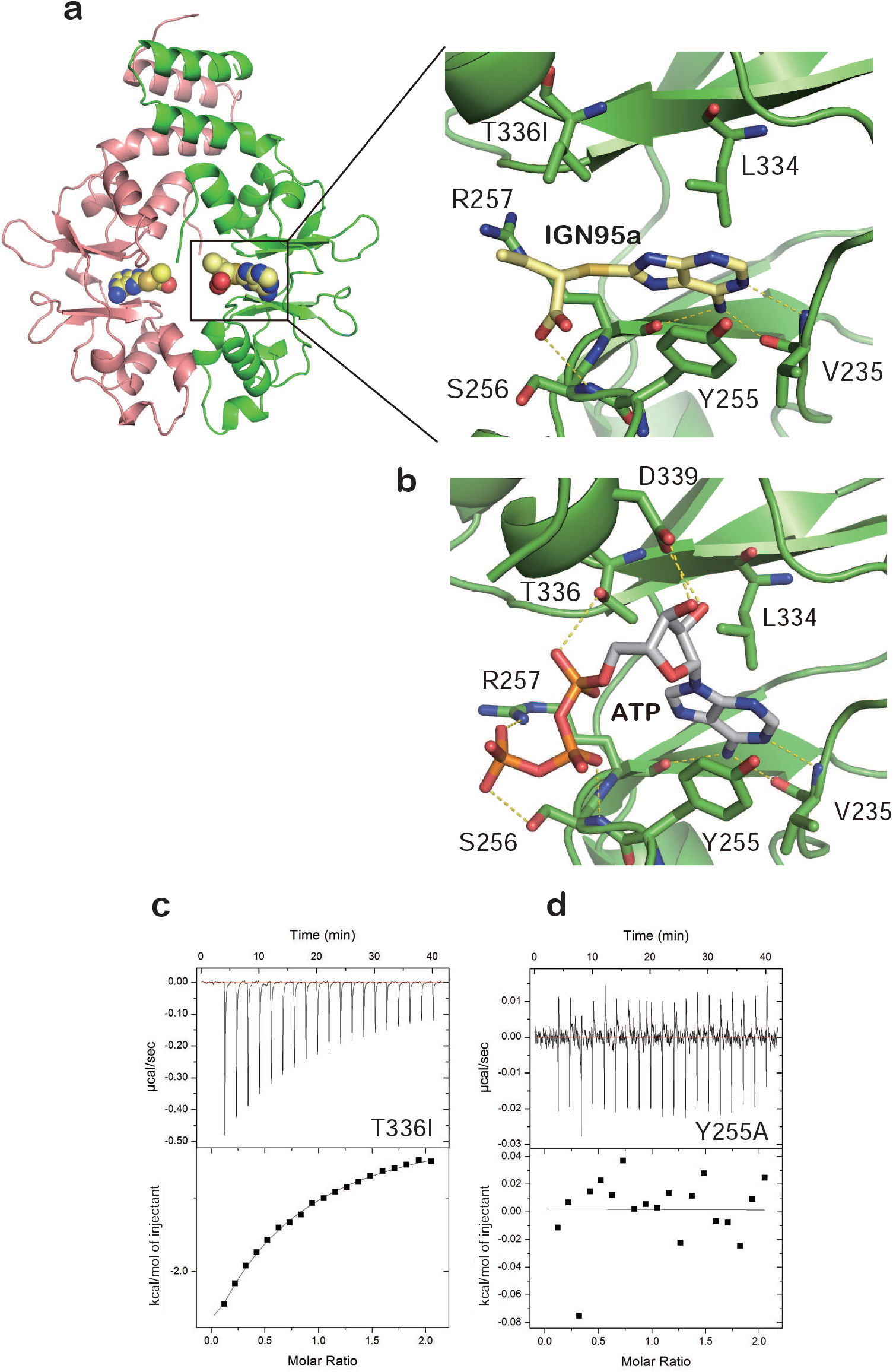
IGN95a-bound structure of the CorC CBS domain. (**a, b**) Close-up views of the IGN95a-binding (**a**) and ATP-binding (**b**) sites in the CorC CBS domain. Ligands and the surrounding residues are shown in stick representation. Dashed lines indicate hydrogen bonds. (**c, d**) ITC data of CorC CBS domain mutants with IGN95a. The raw ITC data and profiles are shown. Measurements were repeated twice, and similar results were obtained.

The adenine ring of IGN95a is recognized by the TpCorC domain (**Fig. 5a**), similar to ATP (**Fig. 5b**). One of the oxygen groups forms an additional hydrogen bond with the main-chain amino group of Ser256 (**Fig. 5a**). In the ATP-bound structure, Thr336, Glu338 and Asp339 form multiple hydrogen bonds with ATP (**Fig. 5b**), but the corresponding residues are not involved in direct interactions with IGN95a (**Fig. 5a**).

The binding pose of IGN95a with CorC in the crystal structure is similar to the docking pose, including the same π-π interaction with Tyr255 (**Supplementary Fig. 2a and 2b**). To examine the stability of the binding of IGN95a to CorC, explicit-solvent molecular dynamics (MD) simulation of the crystal structure of the CorC CBS domain in complex with IGN95a was performed. The representative conformation of the largest clusters of the 300 ns trajectories for IGN95a have similar interactions with the key residues (Val235, Tyr255 and Arg257) in the crystal structure (**Fig. 5a** and **Supplementary Fig. 3a**). Compared to the initial structure, the RMSDs for IGN95a range from 0.2 Å to 1.5 Å (**Supplementary Fig. 3b**), and the key interactions exist during the 300 ns MD simulations (**Supplementary Fig. 3c and 3d**), implying that IGN95a is stable in the binding pocket.

To further verify the structure, we performed a binding assay of the ATP-binding site mutants Y255A (adenine ring) and T336I (ribose) using ITC. According to the structure, mutation at Tyr255 should severely affect IGN95a binding, whereas mutation at Thr336 should have a weaker impact on IGN95a binding (**Fig. 5a**). The T336I mutant of the CBS domain exhibited a *K*_d_ value of 147.3 μM for IGN95a, whereas there were no detectable interactions between the Y255A mutant and IGN95a **(Fig. 5c, d, and Supplementary Table 1**), essentially supporting our structure. Furthermore, since the ethyl group of IGN95a does not exhibit any direct interactions with the CorC CBS domain and is exposed to the exterior of the ATP-binding pocket (**Fig. 5a**), it might be a promising modification target for further optimization of chemical compounds.

### Effect of ATP and IGN95a on the structural equilibrium of the CorC TM domain

To gain insights into the ATP modulation and IGN95a inhibition mechanisms of CorC, we performed biochemical cross-linking experiments using the cross-linking mutant of TpCorC (**Fig. 6**), which we established previously^16^. The TM domain of TpCorC adopts an inward-facing conformation in the presence of Mg^2+^, with an inter-subunit distance of 7.1 Å between the Cβ atoms of Thr106 residues (**Fig. 6a**), which is sufficiently close for chemical cross-linking through Cys residues. We previously generated the cross-linking mutant of TpCorC (T106C/C282A), where Cys282 was also mutated to remove an endogenous cysteine residue. In fact, previous cross-linking experiments with Cu^2+^ phenanthroline showed that the T106C pair of TpCorC formed a disulphide bond in the presence of Mg^2+^, as indicated by a strong band for the dimer on a non-reducing SDS-PAGE gel^16^ (**Fig. 6b**). Furthermore, while the addition of Na^+^ disrupted the cross-linked dimer, the replacement of Na^+^ with K^+^ resulted in bands for both the TpCorC monomer and dimer^16^ (**Fig. 6b**).

**Fig. 6.**
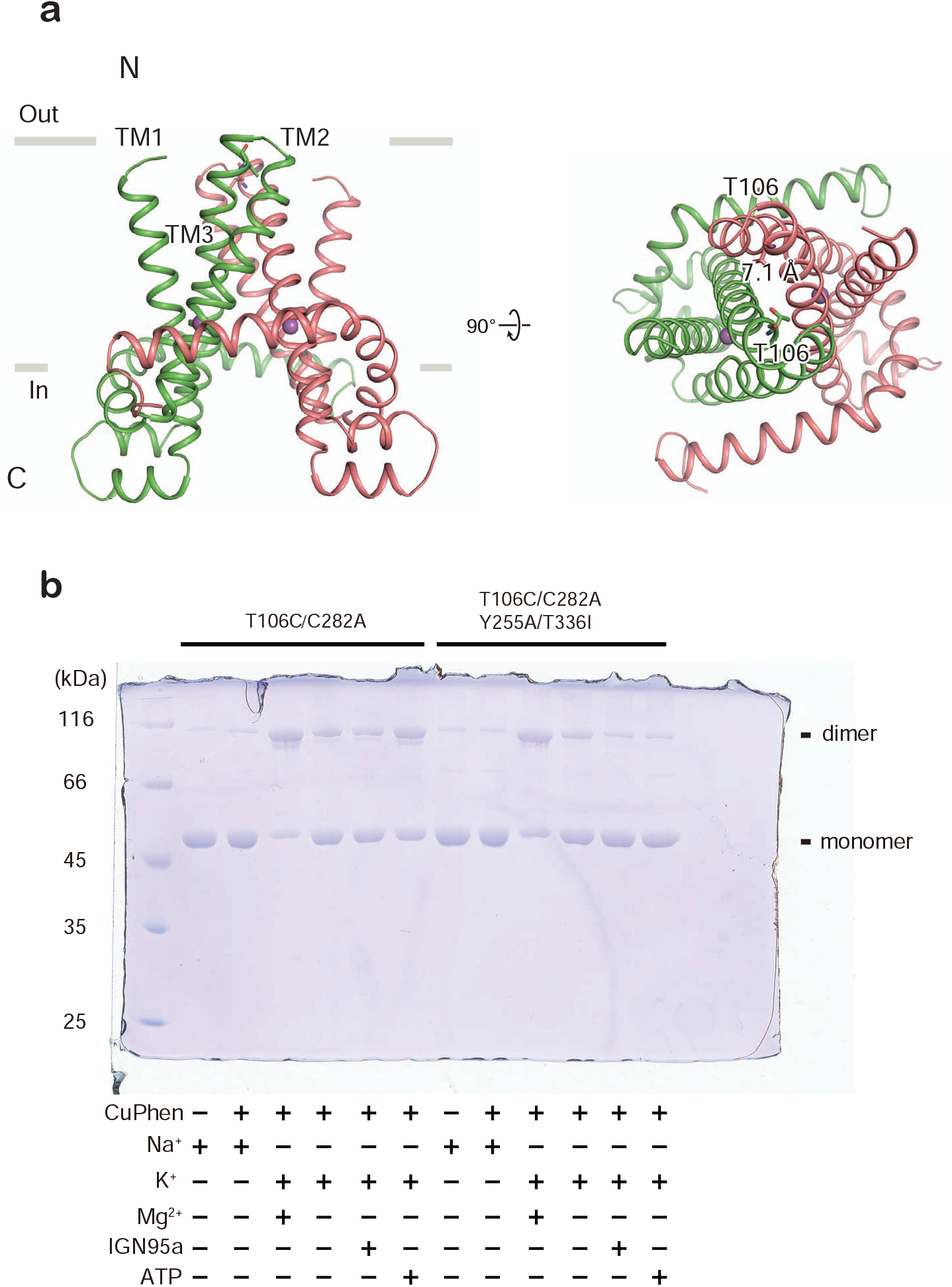
Inter-subunit chemical cross-linking of CorC with ATP and IGN95a. (**a**) Structure of the TpCorC TM domain dimer (PDB ID: 7CFF) in cartoon representation, viewed parallel to the cell membrane (left) and from the periplasmic side (right). The Thr106 residues are depicted in stick representation. Dashed lines show the Cα distances between the Thr106 residues within the dimer. (**b**) Chemical cross-linking experiments of the inter-subunit cross-linking mutant (T106C/C282A) of TpCorC and its ATP-binding site mutant (T106C/T255A/C282A/T336I).

Intriguingly, the addition of ATP to the TpCorC cross-linking mutant in the absence of Na^+^ and Mg^2+^ led to a stronger dimer band than that of the ATP-free sample in the absence of Na^+^ and Mg^2+^ (**Fig. 6b**), whereas we did not see such a shift upon the addition of IGN95a (**Fig. 6b**). Based on this result, we hypothesized that ATP binding to the TpCorC CBS domain affects the conformational equilibrium of the TM domain towards more inward-facing conformations.

To further verify this hypothesis, we generated an ATP-binding site mutant of the TpCorC cross-linking construct (T106C/C282A/Y255A/T336I) for chemical cross-linking experiments (**Fig. 6b**). Mutations in Y255A/T336I are known to abolish the ATP binding activity of TpCorC as well as lower Mg^2+^ export activity^16^. As expected, the addition of ATP to this mutant did not lead to a stronger dimer band than that of the original cross-linking mutant (**Fig. 6b**), further supporting our hypothesis regarding the effect of ATP on TpCorC.

## Discussion

In this work, we performed virtual and *in vitro* screening of CorC by targeting its ATP-binding site and identified a chemical compound, IGN95a, with inhibitory effects on both the ATP binding and Mg^2+^ export activities of CorC (**Fig. 1, 2, 3, 4**). Co-crystallization of the CorC ATP-binding domain with IGN95a and associated MD simulations provided structural insights for the further development and optimization of chemical compounds for the CorC ATP-binding site (**Fig. 5**). Finally, chemical cross-linking experiments indicate that ATP binding to the CorC CBS domain shifts the conformational equilibrium of its TM domain towards more inward-facing conformations, whereas IGN95a, which occupies the ATP-binding site, does not have such an effect (**Fig. 6**). Based on these results, we discuss the mechanisms of action of ATP and IGN95a on CorC (**Fig. 7, 8**)

**Fig. 7.**
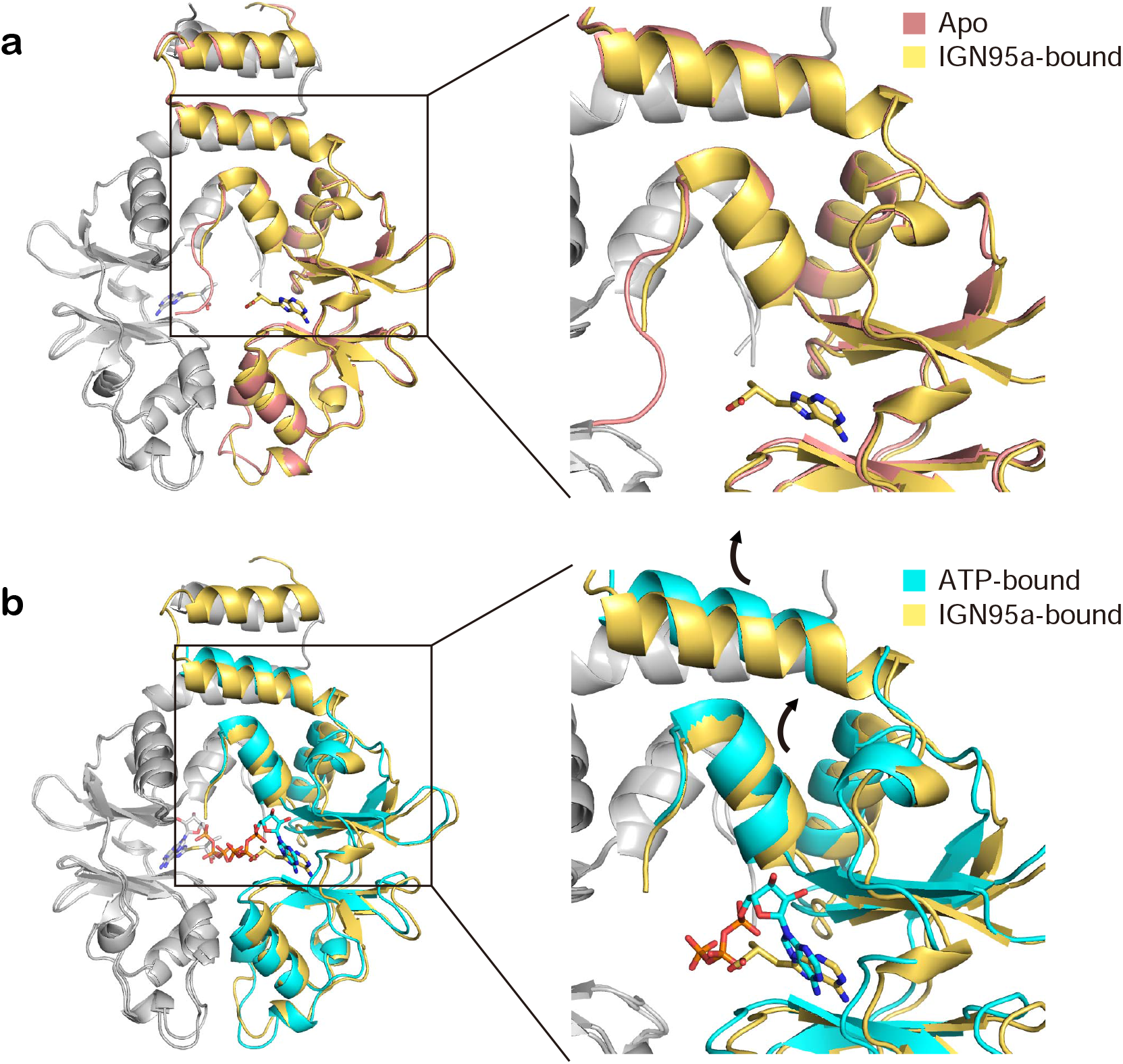
Structural comparisons of the apo, ATP-bound, and IGN95a-bound structures of the CorC CBS domain. (**a**) Superposition of the IGN95a-bound CBS domain structure (yellow) onto the apo structure (red). (**b**) Superposition of the ATP-bound CBS domain structure (cyan) onto the IGN95a-bound structure (yellow). Both subunits in the dimer are shown, and the neighbouring subunit of the dimer is coloured grey. Both the overall structure and close-up view of the region near the exterior of the ATP-binding site are shown. Black arrows indicate the structural changes between two conformations.

**Fig. 8.**
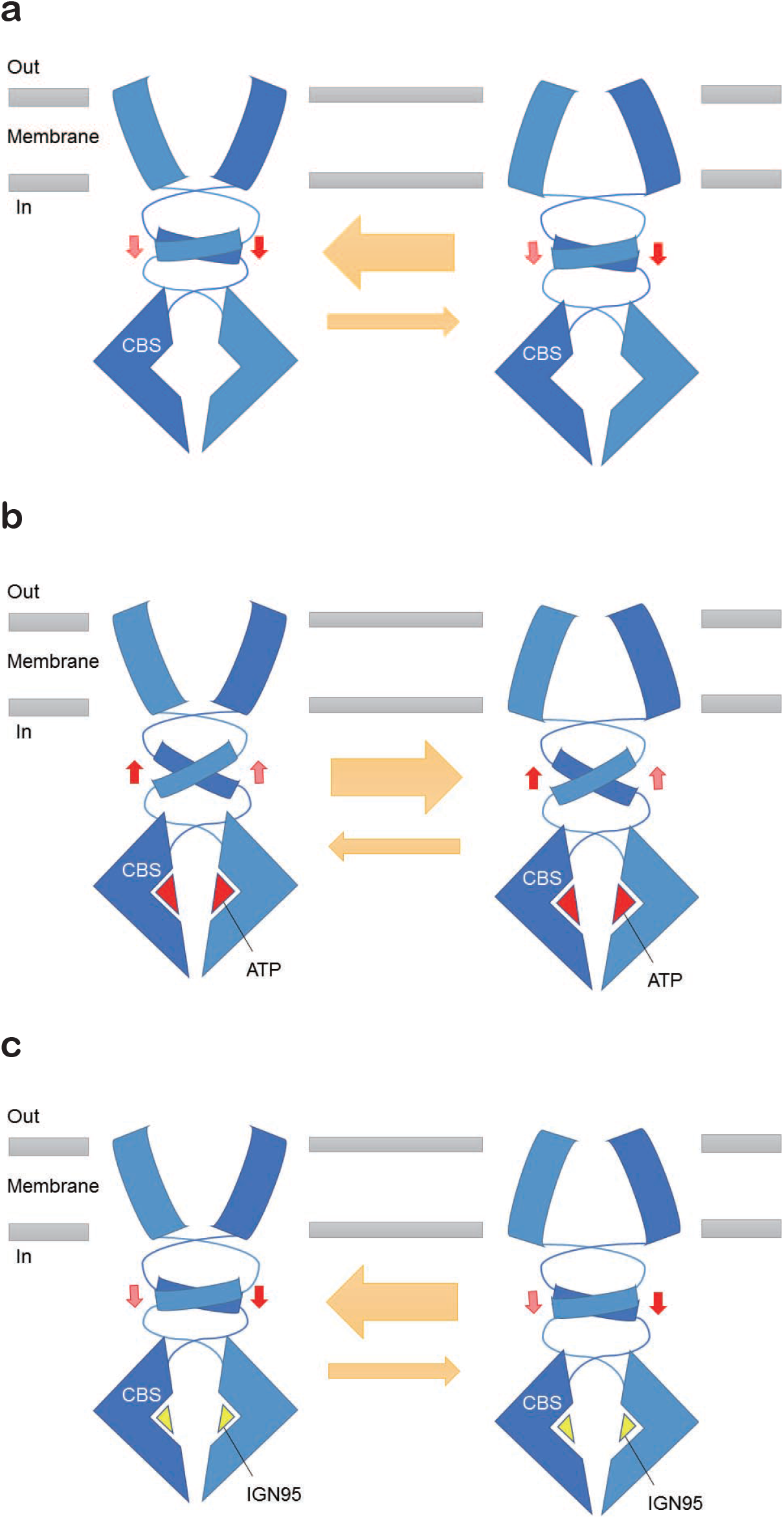
Proposed CorC regulation mechanisms by ATP and IGN95a. (**a, b, c**) Schematic diagrams of the conformational equilibrium of the CorC in the apo (**a**), ATP-bound (**b**), and IGN95a-bound (**c**). ATP binding induces structural changes in the cytoplasmic domain at the interface between the TM and cytoplasmic domains, which shifts the conformational equilibrium of the transmembrane domain more towards the inward-facing state (**a, b**). In contrast, IGN95a binding inhibits the binding of ATP in a competitive manner and inhibits the associated structural changes, which in turn prevents the adjustment of the conformational equilibrium of the TM domain (**c**).

First, a comparison of the apo, IGN95a-bound, and ATP-bound structures of the CBS domain suggests that the IGN95a-bound structure is more similar to the apo structure than to the ATP-bound structure (**Fig. 7).**In the ATP-bound structure, the helix region on the exterior of the ATP-binding site moves slightly outward from the pocket, mainly via its contact with the ribose moiety of ATP (**Fig. 7b**). In contrast, IGN95a binding did not induce such movement (**Fig. 7a)**. Previous chemical cross-linking experiments and MD simulations suggested that this region, undergoing this ATP-dependent structural change, is located at the interface between the TM and CBS domains^16^ (**Supplementary Fig. 4**). Consistent with this, our chemical cross-linking experiments indicated that ATP binding to the CorC CBS domain affects the conformational equilibrium of the TM domain, shifting it towards more inward-facing conformations (**Fig. 6, 8**), which would be more favourable for attraction of intracellular Mg^2+^ to the Mg^2+^-binding pocket in the TM domain. Since the transport activity of CorC is lower in the absence of ATP binding^16^, the ATP-dependent structural change in the CBS domain might be important for CorC to properly maintain its transport activity by adjusting the structural equilibrium of CorC towards a more inward-facing state through the contacts between the TM and CBS domains (**Fig. 8a, b**). Notably, the binding of ATP to the CBS domain of the ClC-1 Cl^−^ channel also affects its transport activity through the domain interface between the TM and CBS domains^28^. In contrast, the binding of IGN95a to the ATP-binding site does not induce the ATP-dependent conformational change of the CBS domain at the interface between the TM and CBS domains (**Fig. 7, 8c, and Supplementary Fig. 4**). Considering the affinity of the CorC CBS domain for ATP^16^ and the cytoplasmic ATP concentration (~mM)^29^, CorC would be mostly in the ATP-bound form *in vivo*. Thus, the addition of IGN95a lowers the transport activity of CorC (**Fig. 4b**), probably because IGN95a binding inhibits the binding of ATP in a competitive manner (**Fig. 5**) and inhibits the associated structural changes (**Fig. 7**), which in turn prevents the adjustment of the conformational equilibrium of the TM domain (**Fig. 6, 8**).

Overall, our results provide not only structural insights for the further design and optimization of chemical compounds targeting the ATP-binding site of CorC but also mechanistic insights into how ATP and chemical compounds modulate the transport activity of CorC.

## Methods

### Protein expression and purification

The CBS domain constructs of the CorC gene from *Thermus parvatiensis* (accession ID: WP_060384576.1) (residues 183-361, C282A) and its mutants were expressed and purified as described previously^16^. In brief, CBS domain constructs with a human rhinovirus (HRV) 3C protease cleavage site and an octa-histidine tag were transformed into the *Escherichia coli* Rosetta (DE3) strain. *E. coli* cells were cultured in LB medium containing 30 μg/ml kanamycin at 37 °C to an OD600 of 0.6, and then, expression was induced with IPTG at a final concentration of 0.5 mM. The *E. coli* cells were then cultured at 37 °C for another 3 hours and harvested by centrifugation (5,000 ×g, 15 minutes). After cell disruption with buffer A [150 mM NaCl, 50 mM Tris (pH 8.0) with 1 mM phenylmethanesulphonyl fluoride (PMSF)], the cell lysate was centrifuged (20,000 ×g, 1 hour). The supernatant was incubated with TALON resin (Takara, Japan) equilibrated with buffer A for 1 hour. The resin was then washed with buffer A containing 10 mM imidazole and eluted with buffer A containing 300 mM imidazole. A His-tagged HRV 3C protease was then added to the eluate to cleave the histidine tag, followed by dialysis with buffer A overnight. The digested sample was reapplied to TALON resin preequilibrated with buffer A. The flow-through fractions from the resin were concentrated by an Amicon Ultra 30K filter (Merck Millipore, USA) and applied to a Superdex 75 Increase 10/300 column (GE Healthcare, USA) in buffer B (100 mM NaCl, 20 mM HEPES (pH 7.5)) for size-exclusion chromatography. The main fractions were collected and concentrated to 10 mg/ml. The CBS domain constructs of human CNNM2 (accession ID: NP_060119.3, residues 433-584) and CNNM4 (accession ID: NP_064569.3, residues 360-511) were similarly expressed and purified.

The C282A and cysteine substitution mutants of TpCorC (residues 23-441) were purified for biochemical cross-linking experiments, as described previously^16^. In addition to the previous purification protocol, to remove endogenous ATP from the purified protein, additional dialysis was performed before solubilization. The collected membrane fraction was transferred into a 10 kDa Spectra/Por 6 dialysis membrane Spectra/Por 6 (Spectrum, USA) and dialyzed against buffer C (1 M NaCl, 50 mM Tris (pH 9.5), 5% glycerol, 2 mM β-ME) for four days by changing buffer C once a day at 4 °C. After 4 days, the membrane was further dialyzed against buffer A containing 2 mM β-ME for 3 hours. In addition, apyrase (Sigma, USA) was added at 0.5 units/ml during cell disruption and membrane solubilization.

### Crystallization

Before crystallization, the crystallization construct of the TpCorC CBS domain (residues 183-361, C282A and T336I) was mixed with IGN95a at a final concentration of 2.5 mM. For crystallization using the vapour diffusion method, 1 μl of the TpCorC CBS domain protein was mixed with 1 μl of reservoir solution (0.1 M KCl, 0.1 M MES (pH 6.0), 32% w/v PEG 400) and stored at 18 °C. Before flash freezing in liquid nitrogen, crystals were incubated with the cryoprotectant solution (0.1 M KCl, 0.1 M MES (pH 6.0), 40% PEG 400, 5 mM IGN95a) for 1 hour and then harvested.

### Data collection and structure determination

X-ray diffraction data sets were collected at the SSRF beamline BL17U1 and the SPring-8 beamline BL32XU and processed using XDS^30^. Notably, at the SPring-8 beamline BL32XU, the X-ray diffraction data sets were collected and processed with the assistance of the automated data collection system ZOO^31^ and the automatic data processing system KAMO^32^.

The structure of the TpCorC CBS domain construct in complex with IGN95a (residues 183-361, C282A and T336I) was determined by molecular replacement with Phaser using the previously determined TpCorC CBS domain structure (PDB ID: 7CFI). The structure was further manually rebuilt with Coot^33^ and refined by Phenix^34^. The Ramachandran plots were calculated using MolProbity^35^. The statistics of data collection and refinement are shown in **Supplementary Table 2**. All structure figures were generated with PyMOL (https://pymol.org/).

### Isothermal titration calorimetry

All ITC measurements were conducted using a MicroCal ITC200 (GE Healthcare, USA) at 25 °C. For ITC experiments, the CorC CBS domain, its mutants and CNNM2 and CNNM4 CBS domain proteins were purified as similarly described above, but buffer D (100 mM KCl, 5 mM MgCl_2_, and 20 mM HEPES (pH 7.5)) was used for size-exclusion chromatography. IGN95a was dissolved in 100% DMSO at a concentration of 100 mM and further diluted to 1 mM with buffer D for ITC experiments. A total of 250 μl of purified proteins (100 μM) was applied to the thermally equilibrated ITC cell. The ligand syringe was filled with 40 μl of IGN95a solution. The ligands were injected 20 times (0.5 μl for injection 1 and 2 μl for injections 2-20), with 120 s intervals between injections. The background data obtained from buffer D containing 1% DMSO were subtracted before data analysis. The data were analysed with Microcal Origin software. Measurements were performed at least twice, and similar results were obtained. The data statistics are summarized in **Supplementary Table 1**.

### Biochemical cross-linking

A total of 4.0 μl of 20 μM TpCorC protein in 20 mM HEPES (pH 7.5) containing 0.03% n-dodecyl-β-D-maltopyranoside (DDM) (Anatrace, USA) and either 150 mM NaCl or 150 mM KCl was mixed with 0.5 μl of 2 mM EDTA, 2 mM EDTA+ 20 mM ATP, 2 mM EDTA+ 20 mM IGN95a, 100 mM MgCl_2_ or Milli-Q water and then incubated for 1 hour at 4 °C. Then, 0.5 μl of the reaction solution (5 mM Cu^2+^ bis-1,10-phenanthroline in a 1:3 molar ratio) or Milli-Q water was added, followed by incubation for 15 minutes at 4 °C. The samples were analysed by non-reducing SDS-PAGE. Experiments were performed at least twice, and similar results were obtained.

### IGN95a

IGN95 (CAS No. 436086-77-0) with a chiral centre was originally purchased from Topscience Co., Ltd. The S-configuration of IGN95 (IGN95a) was separated by WuXi AppTec. The high-performance liquid chromatography (HPLC) purity of IGN95a was greater than 96%. The calculated miLogP was 1.26 (https://www.molinspiration.com/cgi-bin/properties), indicating the cell membrane permeability of IGN95a.

### Molecular docking

A total of 6412 in-house compounds (mostly from commercial databases, e.g., Specs, Maybridge, and ChemDiv, and some synthesized at the Drug Discovery and Design Center, CAS) in pdbqt format were docked to the TpCorC CBS domain by smina^36^, which is a branch of AutoDock Vina^37^ with improved scoring and minimization. Only chain A of the CBS domain (PDB ID: 7CFI) was used for docking, Mg^2+^ ions interacting with the phosphate group of ATP were preserved, and all water molecules were removed. The hydrogens were added to the CBS domain by pdb2pqr (--ff=amber--ffout=amber --chain --with-ph=7)^38^. Then, the model was converted to pdbqt format by the prepare_receptor4.py script in MGLTools version 1.5.6^39^. The geometrical centre of the ATP ligand in the crystal structure was used to define the grid centre, and the grid size was set to 15.0 Å. The random seed was explicitly set to 0. The exhaustiveness of the global search was set to 32, and at most 1 binding mode was generated for each compound. A custom Vina scoring function (**Supplementary Table 3**) was used in this study^40^. The ligand efficiency was calculated as the ratio of the docking score to the number of non-hydrogen atoms in the compound.

### Molecular dynamics simulation of the CBS domain with chemical compounds

Each docking complex or the crystal structure was immersed in a cubic box of TIP3P water that was extended by 9 Å from the solute, and counter ions of Na^+^ or Cl^−^ were added to neutralize the system. The compound and protein were parameterized by the general Amber force field (GAFF)^41^ and Amber ff03 force field^42^, respectively. Ten thousand steps of minimization with constraints (10 kcal/mol/Å^2^) on the heavy atoms of the complex, including 5,000 steps of steepest descent minimization and 5,000 steps of conjugate gradient minimization, were used to optimize each system. Then, each system was heated to 300 K within 0.2 ns, followed by a 0.1 ns equilibration in the NPT ensemble. Finally, 5 ns MD simulations on the docking complex at 300 K were performed for the MM/GBSA calculations. To assess the binding stability of the crystal structure of the TpCorC CBS domain complexed with IGN95a, a 300 ns MD simulation was conducted. Minimization, heating and equilibration were performed with the Sander program in Amber16. The 5 ns or 300 ns production run was performed with *pmemd.cuda.*

### Binding free energy calculation

Based on the first 5 ns of the MD simulation trajectories, the binding free energy (ΔG) was calculated with MM/GBSA^43,44^ according to equation (1):

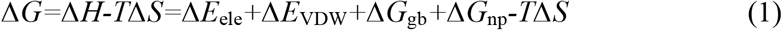

where *ΔE*_ele_ and ΔE_VDW_ refer to the electrostatic and van der Waals energy terms, respectively, and *ΔG*_gb_ and *ΔG*_np_ refer to polar and non-polar solvation free energies, respectively. Conformational entropy (*TΔS*) was calculated by the nmode module in Amber16. The dielectric constants for the solvent and solute were set to 80.0 and 1.0, respectively, and the OBC solvation model (igb = 5 and PBradii = 5)^45^ was used in this study. Other parameters were set to default values.

### Mant-ATP binding assay

The mant-ATP binding assay was conducted in buffer E [50 mM Tris (pH 8.0), 150 mM NaCl and 0.03% DDM]. For saturation binding experiments, a final concentration of 1 μM TpCorC CBS domain protein was mixed with mant-ATP at the indicated concentrations in 96-well plates and incubated in the dark at RT for 1 hour. For ATP-based inhibition experiments, a final concentration of 1 μM TpCorC CBS domain protein and 1 μM mant-ATP were mixed with ATP at the indicated concentrations. To test the effects of chemical compounds, 1 and 5 μM TpCorC CBS protein was preincubated with 0.25 mM and 2 mM chemical compounds, followed by the addition of 1 and 5 μM mant-ATP, as shown in **Fig. 3c** and **Fig. 3d**, respectively, whereas the control samples included the same concentration of DMSO. FRET from endogenous Trp residues to bound mant-fluorophore was measured^26,27^. The fluorescence intensity was measured in an experimental setup with excitation at 280 nm and emission at 450 nm with Cytation 3 (BioTek).

The FRET intensity was calculated by equation (2).

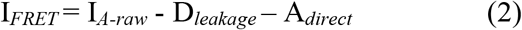

I_*A-raw*_ is the intensity measured at 450 nm with 280 nm excitation, D_*leakage*_ is the leakage of the donor emission into the acceptor wavelength (450 nm) upon donor excitation, and A_*direct*_ is the direct excitation of the acceptor with the donor wavelength (280 nm).

### Mg^2+^ export assay

The Mg^2+^ export assay of TpCorC was performed as described previously^16^. In brief, HEK293-derived cell lines stably expressing TpCorC containing the membrane targeting sequence from human CNNM4 (residues 1-178) were cultured in DMEM (Nissui, Japan) supplemented with 10% FBS, antibiotics and 40 mM Mg^2+^ until used for the Mg^2+^ export assay. The Mg^2+^ export assay using Magnesium Green was performed as described previously^6^. The data are shown as bar graphs of mean relative fluorescent intensities 5 minutes after Mg^2+^ depletion. Relative fluorescence intensities were estimated as the ratio of fluorescence intensity at time 5 minutes to that at time zero. After imaging analyses, HEK293 cells were fixed with PBS containing 3.7% formaldehyde and examined by immunofluorescence microscopy to check protein expression. Statistical analyses were conducted using GraphPad Prism 6 software (GraphPad Software). The data are shown as the mean ± SEM. The *p*-values were calculated by 1-way ANOVA with Holm-Sidak post hoc tests.

## Data availability

The atomic coordinates and structural factors for the structure of the CorC CBS domain in the IGN95a-bound form have been deposited in the Protein Data Bank under accession code 7CFK. All other data are available from the authors upon reasonable request.

## Acknowledgements

We thank the staff from the BL32XU beamline at SPring-8 and from the BL17U1 beamline at Shanghai Synchrotron Radiation Facility (SSRF) for their assistance during data collection. The diffraction experiments were performed at SPring-8 BL32XU (proposal no. 2019A2514) and at SSRF BL17U1 (proposal no. 2018-SSRF-PT-004257). This work was supported by funding provided by the Ministry of Science and Technology of China (National Key R&D Program of China: 2016YFA0502800) to M.H., by funding provided by the National Natural Science Foundation of China (32071234), and by funding provided by the Innovative Research Team of High-level Local Universities in Shanghai and a key laboratory program of the Education Commission of Shanghai Municipality (ZDSYS14005). This work was also supported by funding provided by the Japan Society for the Promotion of Science to H.M. (JP26111007, JP17H04041, and JP20H03515) and to Y.F. (JP20K07312, JP20H05508, and JP17K19396).

## Author contributions

Y.H. and X.T. expressed and purified CorC and its mutants for structural and functional studies and determined the structure. Y.H. and X.T. performed the ITC, and Y.H. and X.T. conducted biochemical cross-linking experiments. Y.F. performed the Mg^2+^ export assay. Z.X., K.M., and W.Z. performed the virtual screening and MD simulations. Y.H. and Y.Z. performed the binding assays for chemical compounds. Y.H., Z.X., K.M., Y.F., H.M., and M.H. wrote the manuscript. Z.X. and M.H. supervised the research. All authors discussed the manuscript.

---

**Supplementary Fig. 1.**
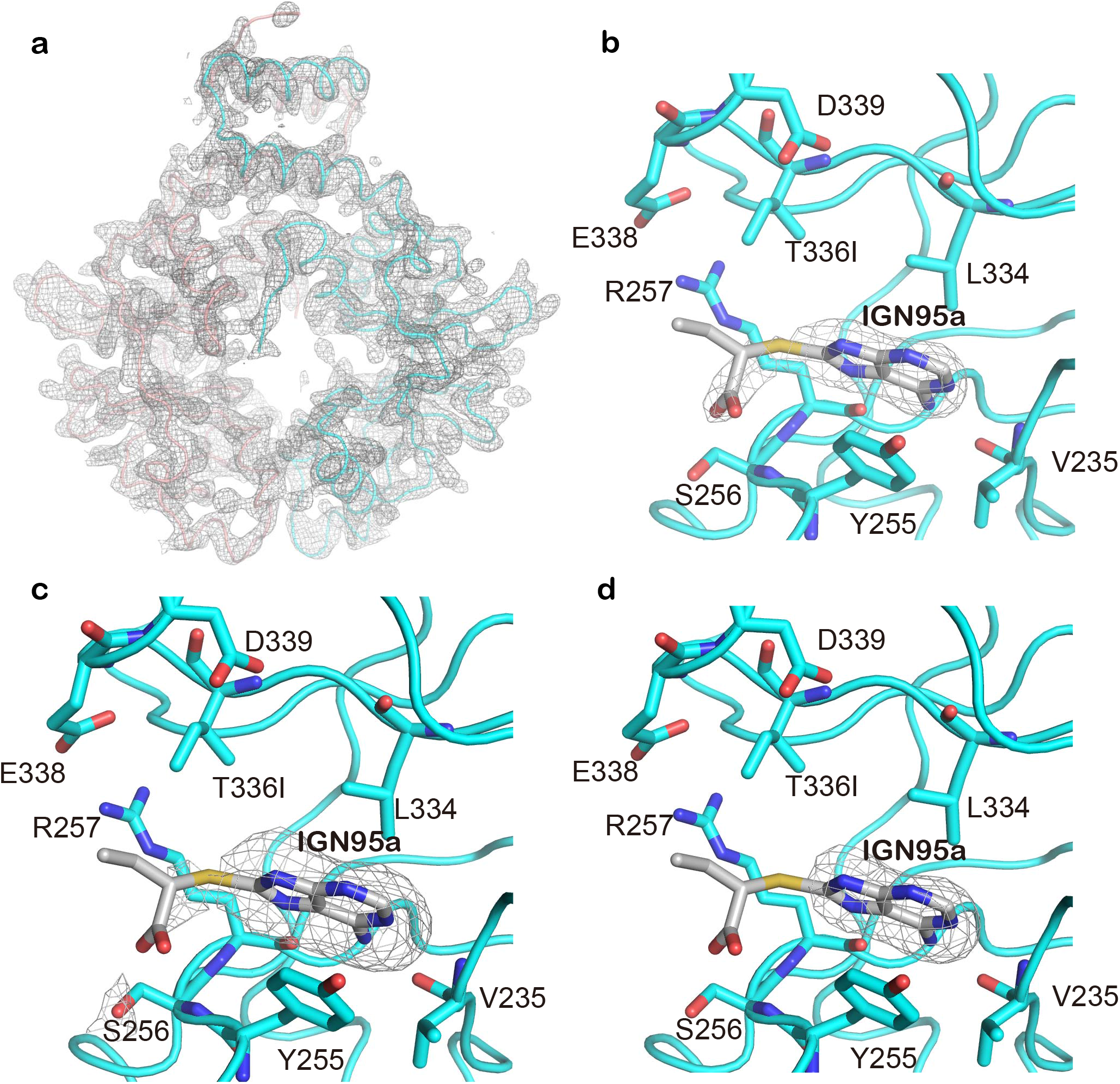
Electron density maps for the IGN95a-bound structure. (**a, b**) The *2F_o_-F_c_* electron density maps for the overall structure of the IGN95a-bound CBS domain (**a**) and for IGN95a (**b**) are contoured at 1 σ, shown in grey mesh. (**c, d**) The *F_o_-F_c_* omit maps for IGN95a are contoured at 1.8 σ (**c**) and 3.0 σ (**d**) and are shown in grey mesh.

**Supplementary Fig. 2.**
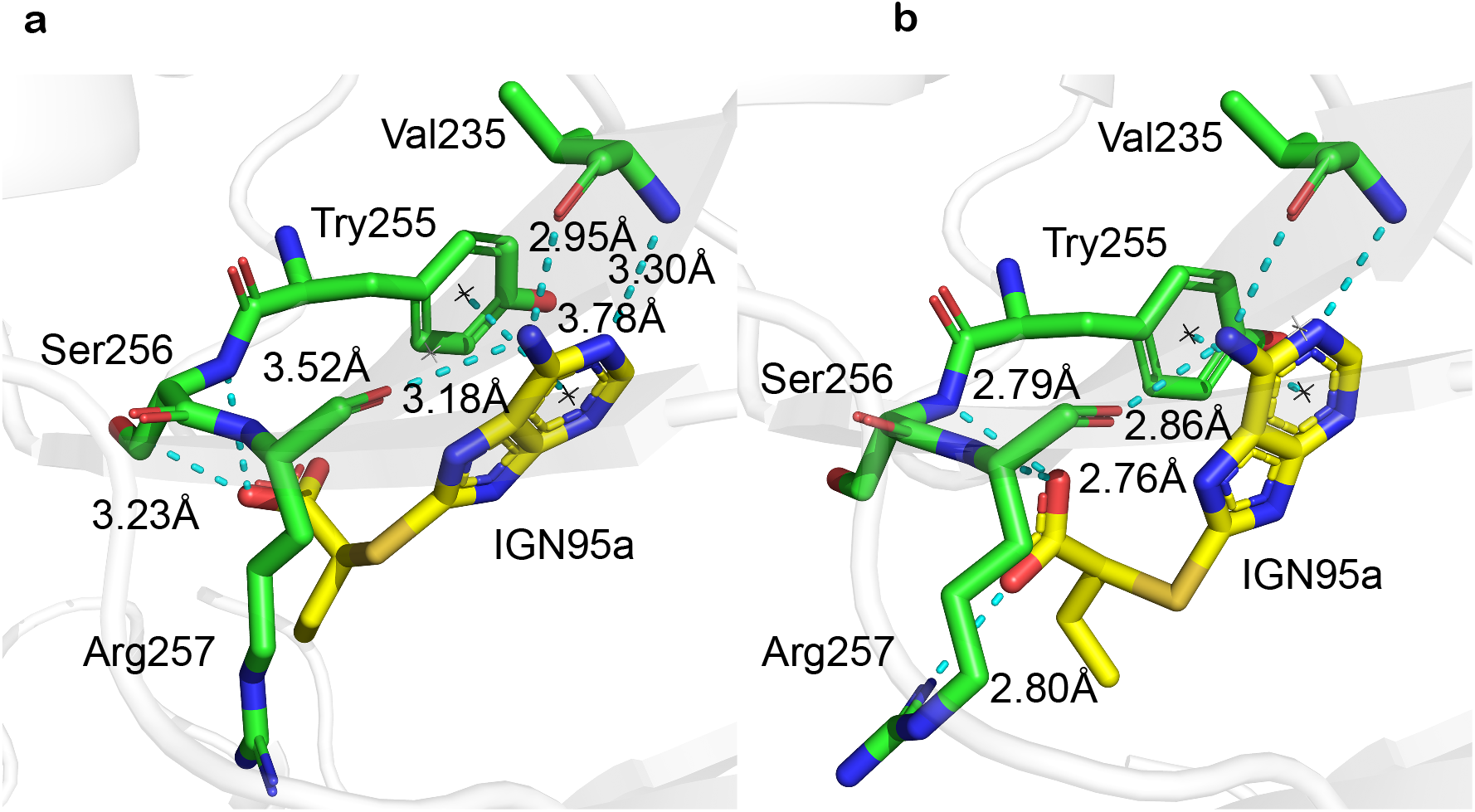
Comparison of the IGN-bound CorC CBS domain structure and the corresponding docking model from virtual screening. (**a**) The binding pose of IGN95a with TpCorC in the crystal structure. (**b**) The docking pose of IGN95a against TpCorC. The proteins are shown in grey cartoon, the ligands in yellow sticks, and key residues in green sticks. The non-covalent interactions shown as cyan dotted lines with the distance in Å.

**Supplementary Fig. 3.**
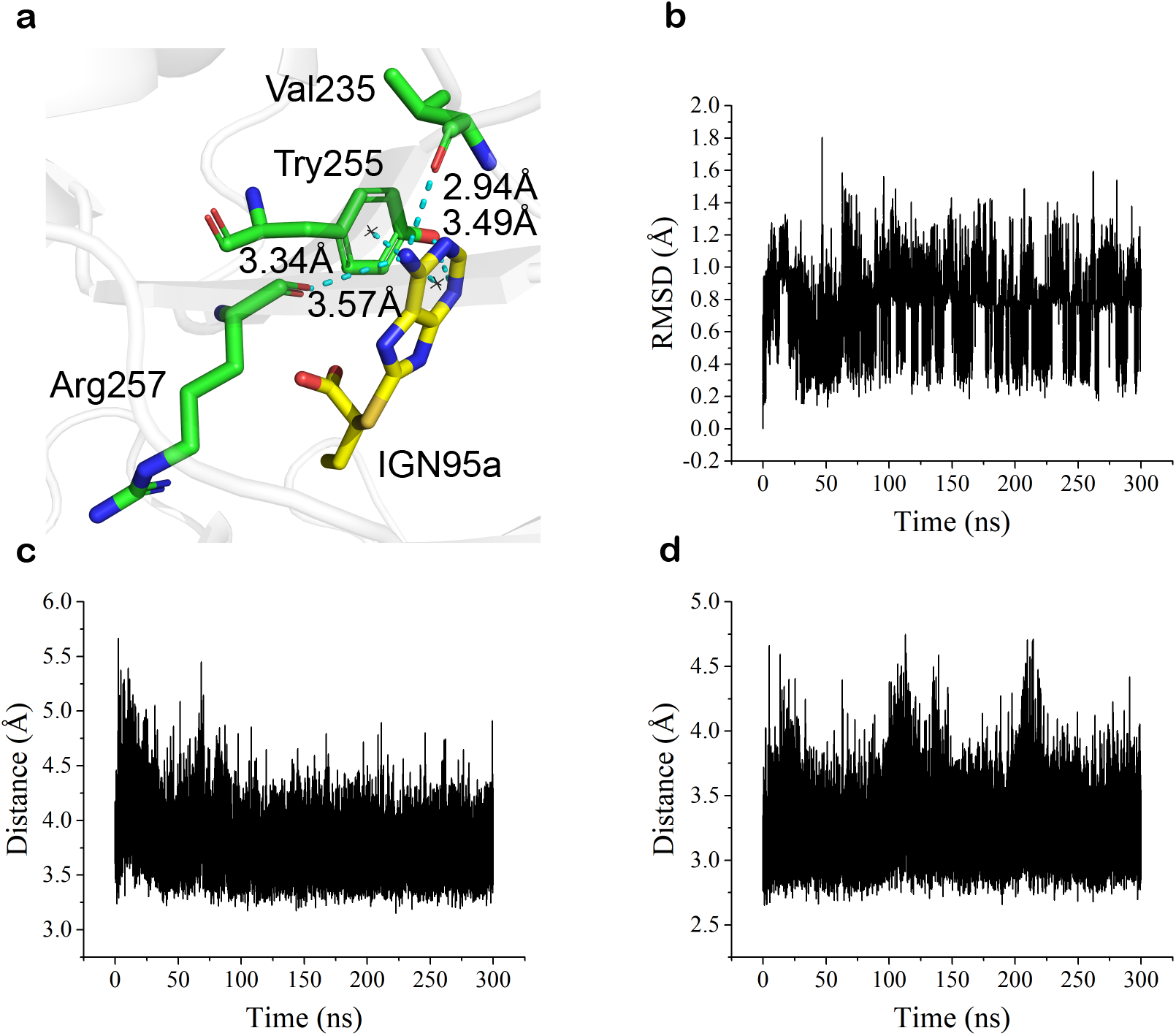
MD simulations. (**a**) The interactions between IGN95a (green sticks) and protein residues (yellow sticks) of the representative conformation during 300 ns trajectories. (**b**) The RMSDs to the initial structure of IGN95a during 300 ns MD simulations. (**c**) Distances between the benzene ring of IGN95a and the indole ring of Tyr255 during 300 ns MD simulations. (**d**) Distances between the amino nitrogen of IGN95a and the carbonyl oxygen of Val235 during 300 ns MD simulations.

**Supplementary Fig. 4.**
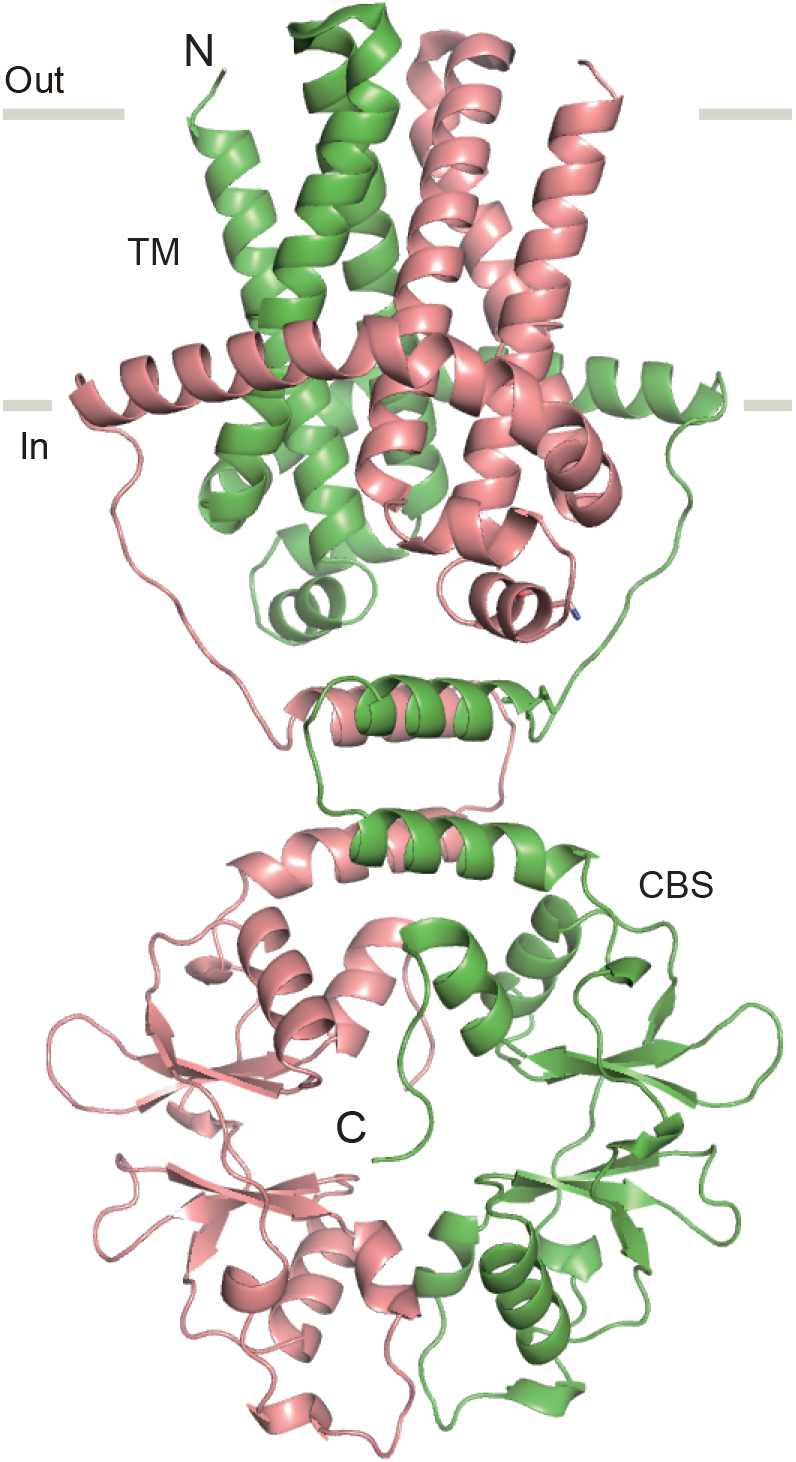
Structural model of TpCorC TM-CBS. Viewed parallel to the membrane. The model was previously reported based on chemical cross-linking experiments and MD simulations^16^. The two subunits are coloured red and green.

**Supplementary Table 1.** ITC statistics.

| **TpCorC CBS**<br>**with IGN95a** | **n** | **$\Delta H^\circ$**<br>**(kcal/mol)** | **$-T\Delta S^\circ$**<br>**(kcal/mol)** | **$\Delta G^\circ$**<br>**(kcal/mol)** | **$K_D$ (μM)** |
| --- | --- | --- | --- | --- | --- |
| WT | $0.23 \pm 0.01$ | $-6.7 \pm 1.2$ | 0.8 | $-5.9 \pm 1.2$ | $46.95 \pm 4.23$ |
| T336I | $0.25 \pm 0.10$ | $-18.2 \pm 8.3$ | 12.9 | $-5.3 \pm 8.3$ | $147.28 \pm 18.35$ |
| Y255A |  |  |  |  | ND |
| **CNNM CBS**<br>**with IGN95a** | **n** | **$\Delta H^\circ$**<br>**(kcal/mol)** | **$-T\Delta S^\circ$**<br>**(kcal/mol)** | **$\Delta G^\circ$**<br>**(kcal/mol)** | **$K_D$ (μM)** |
| HsCNNM2 |  |  |  |  | >1000 |
| HsCNNM4 |  |  |  |  | > 500 |

**Supplementary Table 2.**
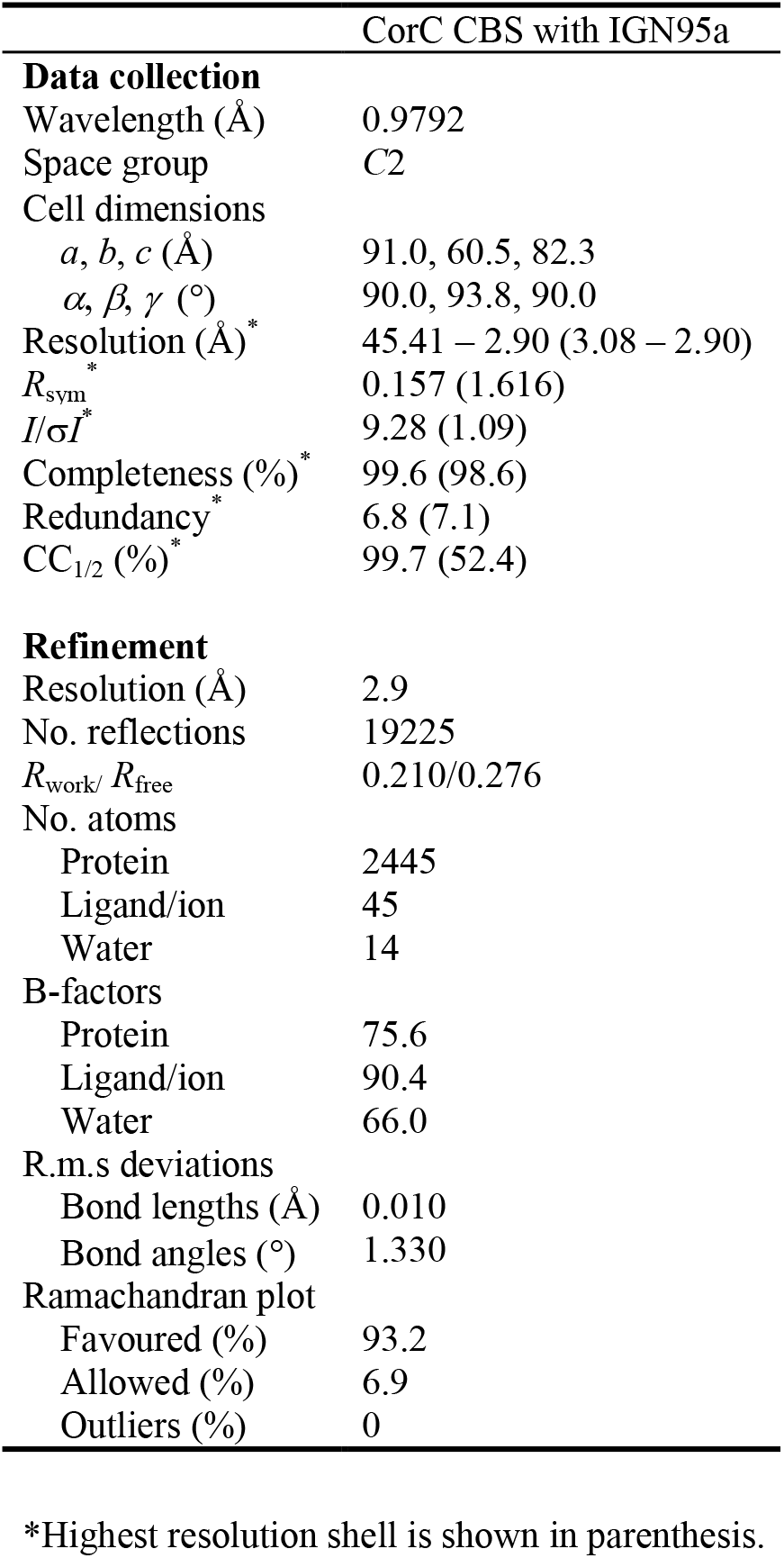
X-ray Data collection and refinement statistics

**Supplementary Table 3.** Customized parameters for molecular docking.

| Parameters | Weights |
| --- | --- |
| gauss (o=0, w=0.5, _c=8) | -0.035579 |
| gauss (o=3, _w=2, _c=8) | -0.005156 |
| repulsion (o=0, _c=8) | 0.26360521 |
| hydrophobic (g=0.5, _b=1.5, _c=8) | -0.035069 |
| non_dir_h_bond (g=-0.7, _b=0, _c=8) | -0.587439 |
| num_tors_div | 1.923 |
| atom_type_gaussian(t1=Magnesium,t2=OxygenXSAcceptor,o=0, w=3, c=8) | -0.3 |

